# Development of a High-throughput Morphological Assay for Evaluating Mesenchymal Stromal Cell-derived Extracellular Vesicle Modulation of Brain Pericyte Secretory Phenotype

**DOI:** 10.1101/2025.01.15.633196

**Authors:** Courtney E. Campagna, Andrew M. Larey, Kanupriya R. Daga, Morgan Roos, Sneha Ghosh, Neil Grimsey, Jin Han, Ross A. Marklein

## Abstract

Mesenchymal stromal cell-derived extracellular vesicles (MSC-EVs) are a promising therapeutic tool for treating many neurodegenerative diseases. Neuroinflammation plays a major role in many of these conditions through an orchestration of interdependent processes that lead to the breakdown of the blood-brain barrier (BBB), infiltration of immune cells and neuronal death. MSC-EVs have shown preliminary evidence of modulating neuroinflammation, but their mechanisms of action are still unknown. Therefore, we explored the potential of MSC-EVs in modulating brain pericytes, a cell type that plays a critical role in BBB maintenance but has not been investigated as a therapeutic target for MSC-EVs. Brain pericytes are multifaceted cells that can modulate neuroinflammation through their involvement in BBB homeostasis, as well as the innate and adaptive immune response. Pericyte morphology has been shown to change in response to inflammatory stimuli *in vivo*, hence, we used this behavior to develop a quantitative morphological profiling approach to assess the immunomodulatory function of MSC-EVs in a high-throughput, low-cost manner. Using this assay, we were able to demonstrate that MSC-EVs manufactured under various conditions (2D, 3D, and in response to cytokine priming) could induce distinct pericyte morphological responses indicative of changes in secretion of chemokines and cytokines relevant to neuroinflammation.

## INTRODUCTION

Neurodegenerative diseases lead to the loss of cognitive and functional abilities due to neuronal death. Despite varying underlying causes, neuroinflammation is a common factor across many of these conditions, posing a major challenge for therapy development^1,2^. Neuroinflammation is an immune response within the central nervous system (CNS), involving interactions between leukocytes, microglia, astrocytes, pericytes, glial cells, and their associated signaling molecules^3^. In homeostatic conditions, neuroinflammation can serve a beneficial role to protect and promote normal brain function by regulating the recognition, trafficking, and elimination of pathogens and cell debris^4^. However, when neuroinflammation becomes chronic or dysregulated, it can result in tissue damage and further progressing disease pathology^3^. Some detrimental aspects of neuroinflammation include the release of pro-inflammatory cytokines and chemokines, activation of immune cells, the breakdown of the vasculature, and neuronal death^4^. These adverse effects are often associated with the breakdown of the blood-brain barrier (BBB), the interface between the blood and the brain parenchyma that maintains the CNS stability. The BBB, composed of pericytes, astrocytes, and brain endothelial cells (BMECs), plays a key role in maintaining CNS homeostasis by regulating the passage of molecules across the blood-brain interface^5^. Therefore, preserving BBB integrity, and regulating neuroinflammation necessitates an immunomodulatory, multi-cellular approach to prevent further disease progression.

Mesenchymal stromal cells (MSCs) have been explored as a therapeutic for inflammatory diseases due to their immunomodulatory behavior. In terms of neuroinflammation, MSCs have shown promise in treating neurodegenerative diseases such as traumatic brain injury (TBI) and stroke in animal models^6,7^. However, due to the tightly regulated nature of the BBB, MSCs cannot readily enter the brain parenchyma when administered intravenously with most of injected MSCs becoming trapped in the lung^8,9^. Additionally, local intracerebral injection of MSCs comes with inherent risks of damaging adjacent healthy tissues, even with efficient delivery of the cells^8^. Moreover, heterogeneity of MSCs has also posed challenges in comprehending both their mechanism of action and any associated safety concerns.

To address some of the challenges associated with MSCs, MSC-derived extracellular vesicles (MSC-EVs) have garnered significant interest in treating neuroinflammation. MSC-EVs are lipid-membrane-bound structures secreted by MSCs that contain bioactive molecules such as proteins, metabolites, lipids, and nucleic acids^8^. MSC-EVs are non-replicative and small sized (∼50-1000s nm in diameter), posing minimal inherent risks of tumorigenicity or thrombosis^10,11^, and can readily cross the BBB^12^. Importantly, MSC-EV cargo, which includes tetraspanins, receptors, integrins, lipids, and miRNA, exhibits immunomodulatory functions similar to those of MSCs^13,14^. Moreover, MSC-EV immunomodulatory function can also be enhanced by exposing the source MSCs to inflammatory-relevant signals (e.g. cytokines, hypoxia or low pH) for a short period of time (1-3 days), termed ‘priming’^13^. For example, MSCs primed with reduced serum and 1% oxygen for 48 hours enhanced secretion of proteins with mitogenic and neurotrophic functions^15^. Another study showed that cytokine-primed MSCs cultured in hypoxic condition further impacted lipid composition of MSC-EVs and enhanced their functionality to modulate microglia morphology^2^. While these studies provided promising preliminary evidence on the effects of manufacturing (e.g. priming) on MSC-EVs in the context of neuroinflammation, there is currently no standardized method for evaluating their immunomodulatory function. The current recommendations of the International Society of Extracellular Vesicles (ISEV) focus primarily on EV identification with no specific recommendations regarding functionality or immunomodulatory activity^16^. Although some studies have employed multi-omics approaches and wound healing assays both *in vitro* and *in vivo* to assess EV function^17–20^, these methods are often costly, low-throughput, difficult to standardize, and not specifically tailored to neuroinflammation.

Pericytes are multipotent cells^21,22^ located throughout the basement membrane of the vasculature. They are heterogeneous in origin, tissue distribution, morphology, and play a critical role in the progression of neuroinflammation due to their regulation of the BBB, as well as proximity^23^. Pericytes assist in the regulation of vasculature including angiogenesis and capillary blood flow, as well as the regeneration of function and structure in damaged parts of the CNS^23^. Additionally, they regulate various aspects of the immune response, such as leukocyte extravasation, inflammation-induced BBB disruption, propagation of peripheral and central inflammation, and polarization of the inflammatory cells in the BBB and the brain parenchyma^24^. These diverse functions make pericytes a strong candidate target for studying neuroinflammation and for screening potential therapies. However, while they have been more actively studied as a target for drug therapies^25^, the effects of MSC-EVs on this versatile cell type remain largely unknown. *In vivo*, pericytes undergo morphological changes in response to inflammation, becoming activated and often migrating away from the sites of chronic inflammation. For instance, following penetrating cortical injury in humans, several morphological types of pericytes were observed near injury site, migrating away from blood vessels^26^. Interestingly, MSCs also respond to inflammatory stimuli (in the form of cytokine priming) through morphological changes similar to pericytes. We have previously shown this morphological shift can predict a given MSC batch’s immunomodulatory function, and help screen priming conditions to optimize MSC manufacturing^27,28^. For example, a morphological screening of MSCs primed with various combinations of Interferon-gamma (IFN-γ) and Tumor Necrosis Factor-alpha (TNF-α) identified optimal IFN-γ/TNF-α priming conditions that enhanced MSC function in terms of T cell suppression^28^.

Previous works suggested pericytes and MSCs exhibit phenotypic similarities, such as cellular morphology and cell surface molecules^29,30^. Hence, we hypothesize that a similar morphological profiling approach with pericytes could serve as an indicator of their response to inflammation and immunomodulatory effects of MSC-EV^31^. Here, we developed a high content imaging-based morphological assay on pericytes to assess the effect of MSC-EVs generated under varying manufacturing conditions (e.g., flask vs. bioreactor, priming, microcarrier density). TNF-α was used as an inflammatory stimuli to activate pericytes, as it plays a key role in neuroinflammation and many neurodegenerative diseases, including Alzheimer’s Disease, Ischemic stroke, Traumatic Brain Injury^32–34^. We present a comprehensive single-cell morphological profiling of pericyte responses to TNF-α and MSC-EVs from various manufacturing conditions. We further explore the relationship between these pericyte morphological responses and secretome profiles of cytokines and chemokines.

## MATERIALS AND METHODS

### Pericyte Expansion and Cryopreservation

Human Brain Pericytes (ScienCell) were expanded in Pericyte Complete Medium (ScienCell) on T175 flasks at 3500 cells/cm^2^ coated with 1% (w/w) Poly-d-lysine (PDL) (Sigma) for four passages with TrypLE (Gibco) used to harvest cells at the end of each passage upon reaching 80-90% confluency. At passage 4, pericytes were cryopreserved in pericyte media containing 10% DMSO.

### MSC-EV Manufacturing

Human bone marrow derived MSC line RB71 was expanded according to the RoosterBio protocol previously described^28^ and then cryopreserved at passage 2. One frozen vial containing 10^6^ MSCs was seeded into T225 flasks at a seeding density of 4,444 cells/cm^2^ in RoosterNourish-MSC-XF and cultured to 80% confluency. For flask MSC-EV manufacturing, the MSCs were passaged into T175 flasks at a seeding density of 714 cells/cm^2^. For the bioreactor group, MSCs were passaged into 0.5L spinning wheel bioreactors (PBS Biotech) with 0.4g Corning Synthemax II polystyrene microcarriers. These MSCs were expanded in their respective vessels following protocols from PBS Biotech and RoosterBio. The MSCs were then primed in serum-free RoosterCollect-EV medium with control and priming conditions. The MSC-EVs were collected from the supernatant using an adapted 2-step ultracentrifuge protocol^35^. The supernatant was first filtered through a 0.2µm filter and then centrifuged at 133,900 xg (Soorvall WX ultracentrifuge, ThermoFisher; Fiberlite F37L-8×100 Fixed-Angle Rotor, ThermoFisher, k factor=224; PC Bottle Assembly 70mL, ThermoFisher) for 1 hour at 4°C. The MSC-EV pellet was then resuspended in cold PBS-/- and then centrifuged again in micro-ultracentrifuge tubes (PC Thickwall 4mL, ThermoFisher) at 140,000xg (Sorvall MX 150+ Micro-Ultracentrifuge, ThermoFisher; S110-AT Fixed-Angle Rotor, ThermoFisher, k factor=76) for 1 hour at 4°C. The EV pellets were then resuspended at a 37.5X concentration in cold PBS-/-. The prepared EVs were then stored at −80°C until used.

### Pericyte Morphological Assay

A T175 flask was coated with 1% (w/w) poly-d-lysine (PDL) for 24 hours at 4°C. After, the flask was washed with PBS three times to get rid of excess PDL and 35 mL of pericyte medium (Sciencell) was added. 10^6^ previously frozen human brain pericytes (Sciencell) were thawed and seeded on the coated T175 flask. A flat-bottomed 96-well plate (Corning, Cat # 3599) was coated with 1% (w/w) poly-d-lysine (PDL) for 24 hours at 4°C. Once the pericytes reached 80-90% confluency (3 days) they were harvested using TrypLE (Gibco) and were seeded at 2,500 cells/cm^2^ for 24 hours in 100 µL of pericyte complete medium in the prepared 96-well plate. After 24 hours, 50% of the media (50 µL) was aspirated from each well. Then 50 µL of pericyte medium only (‘unstimulated’) or 50 µL of pericyte medium containing 50 ng/mL TNF-α (Gibco, PHC3015) (‘stimulated’) were added to appropriate wells and pericytes incubated for an additional 24 hours. For MSC-EV treated pericyte groups, MSC-EVs were added concurrently with 100 ng/mL of TNF-α (50 ng/mL final concentration due to half media change) in pericyte media to the appropriate wells. We fixed samples using 4% paraformaldehyde (Electron Microscopy Sciences) and stained with Hoechst [10µg/mL] (Invitrogen) and Fluorescein maleimide [20µM] (ThermoFisher) in PBS-/- (Gibco) for nuclear and cytoplasm morphology, respectively as done previously^27^. Pericytes were imaged using Cytation 5 High Content Imaging system (Agilent) and a Ti-Eclipse (Nikon) with the imaging system indicated in each figure legend. We imaged approximately 50% of every well using a 6×6 montage at 10X magnification. These images were processed on a single-cell basis for over 96 different nuclear and cytoplasmic morphological features using CellProfiler pipeline^36^ (Supplementary File 1), however, we focused primarily on the 21 features^27^. A table summarizing and defining all morphological features quantified in this study can be found in Supplementary Table 1.

### Pericyte Secretome

Pericytes were cultured on a PDL coated 96-well flat-bottomed plate at a density of 22,580 cells/cm^2^ for 24 hours in pericyte media. After 24 hours, 50% of the media (50 µL) was aspirated from each well. Then 50 µL of pericyte medium only (‘unstimulated’) or 50 µL of pericyte medium containing 50 ng/mL TNF-α (Gibco, PHC3015) (‘stimulated’) were added to appropriate wells and pericytes incubated for an additional 24 hours. For MSC-EV treated pericyte groups, MSC-EVs were added concurrently with 100 ng/mL of TNF-α (50 ng/mL final concentration due to half media change) in pericyte media to the appropriate wells. Pericyte conditioned medium was collected from n=3 wells per experimental group and stored at −80°C. Frozen CM samples were shipped on dry ice to RayBiotech (Norcross, GA) for secretome analysis using the 200plex quantibody Array (Q4000). Differentially secreted proteins of interest were analyzed using STRING database (https://string-db.org/) to identify significantly enriched pathways.

### Data Analysis and Statistics

Singe cell morphological data is presented on a median-per-well basis and summarizes data from at least 300 cells per well. All data and statistical analyses were performed using GraphPad Prism v10 with specific statistical tests performed are detailed in each figure legend.

## RESULTS

### TNF-α-stimulated pericytes exhibit increased size and morphological complexity

Human brain pericytes were expanded until passage 4, seeded into a 96-well plate coated with Poly-D-lysine (PDL) and treated with or without [50 ng/mL] TNF-α (**Figure 1A**). Qualitatively, we observed that TNF-α-stimulated pericyte (TNF-α) morphology was different from that of unstimulated pericytes (CTL: **Figure 1B**) as reflected in the significant and quantifiable change of multiple morphological features including perimeter, major axis length, compactness, form factor and aspect ratio (**Figure 1C**). The stimulated pericytes increased significantly in perimeter and major axis length, illustrating an overall increase in size. A notable increase in aspect ratio also indicated a more elongated morphology upon stimulation, which coincided with the observed increase in major axis length. In addition, the stimulated pericytes decreased in form factor while increasing in compactness, which indicated an overall increase in complexity of morphology. Significant quantitative and qualitative changes in pericyte shape were consistently observed across six independent experiments, suggesting a robust morphological response to TNF-α stimulation.

**Figure 1.**
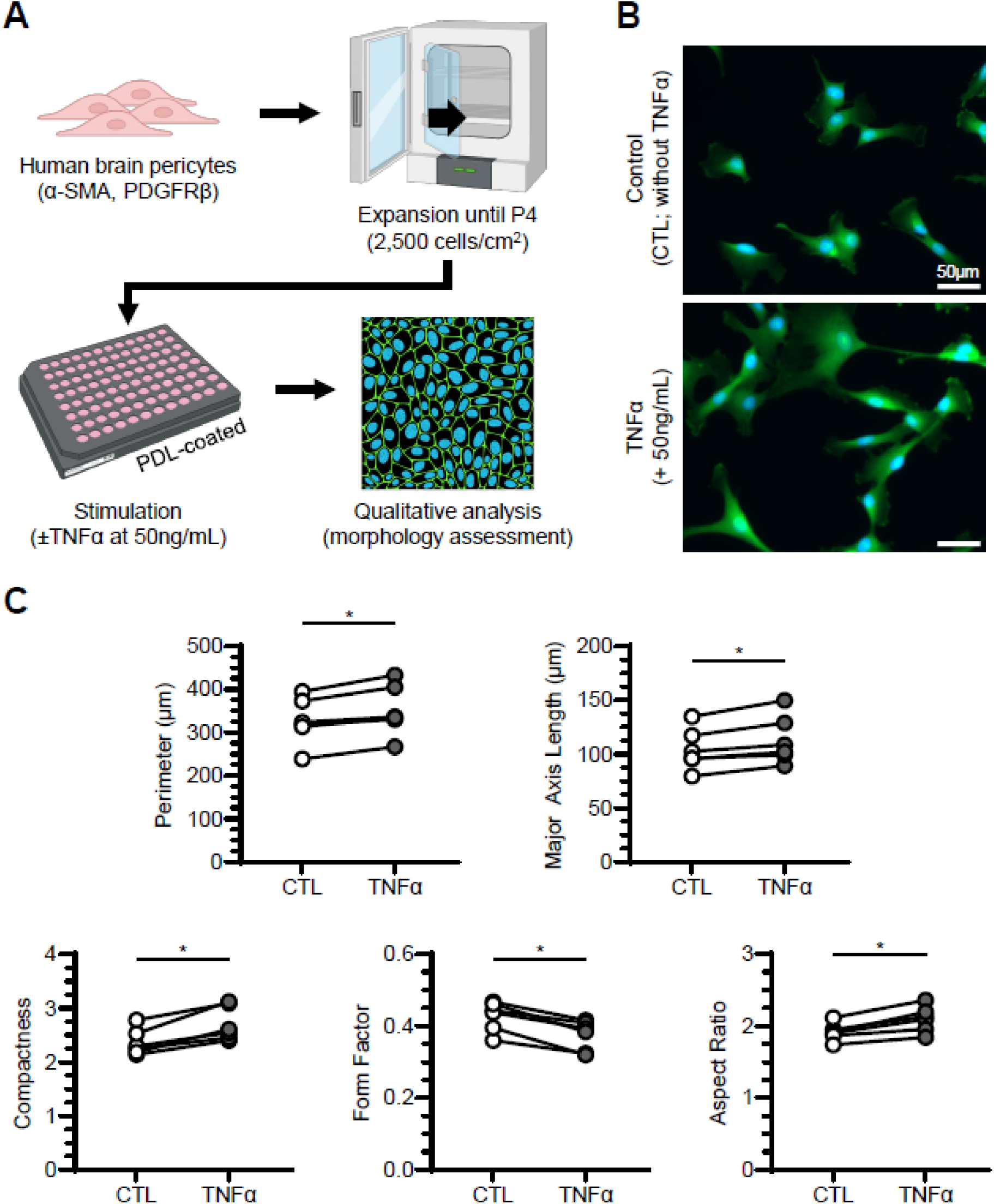
Pericytes become more complex and elongated with TNF-α stimulation. **A**. Schematic illustration of pericyte culture and their morphological assessment before and after TNF-α stimulation. **B.** Representative images of unstimulated pericytes (CTL; top) and pericytes stimulated with 50ng/mL of TNF-α (TNF-α; bottom). Pericytes were stained with Hoechst (blue) and fluorescein-maleimide (green) to indicate nuclei and cytoplasm, respectively. **C.** Quantification of morphological parameters between unstimulated (open circles) and stimulated pericytes (filled circles). For each morphological feature, each data point represents average of six replicate wells for given experiment (n=6 independent experiments). *p<0.05.

### Immunological stimulus promotes pro-inflammatory secretory phenotype in pericytes

To evaluate changes in protein secretory phenotype in addition to the morphological changes after immunological stimulation, we further performed quantitative secretome profiling on unstimulated and stimulated pericytes (**Figure 2**). Pericytes were cultured on a 96-well plate with or without TNF-α stimulation, after which the supernatant was collected and analyzed using the Human Cytokine Array Q4000 (RayBiotech). 81 total proteins were detected as secreted by pericytes at levels above the pericyte medium-only control and were used for further analysis (**Supplementary Table 2**). Hierarchical clustering, which compared unstimulated to TNF-α-stimulated pericytes with three replicates in each group, illustrated a distinct secretion profile from pericytes after stimulation (**Figure 2A**). Among 81 proteins, 18 proteins exhibited significant changes upon TNF-α stimulation, and further evaluation of these proteins using STRING analysis highlighted several proteins that were highly connected (based on node degree), such as IL-6, ICAM1, and CXCL10 (**Figure 2B**). Proteins ordered by node degree, which indicates the number of connections in **Figure 2B**, is shown in **Supplementary Table 3**. Among the 18 differentially expressed proteins, 16 were increased and 2 were decreased after TNF-α stimulation (**Figure 2C**). Notably, the increased proteins included ICAM1 and VCAM1, which are relevant to pericyte physiology and morphology through mechanisms of adhesion, signaling, and response to inflammatory stimuli—indicating that immunological stimulation may alter cellular morphology and their physical function. Other increased proteins included pro-inflammatory chemokines CCL5, CSF3, CCL23, and TEK, which were below the threshold of detection in unstimulated groups but dramatically increased upon stimulation.

**Figure 2.**
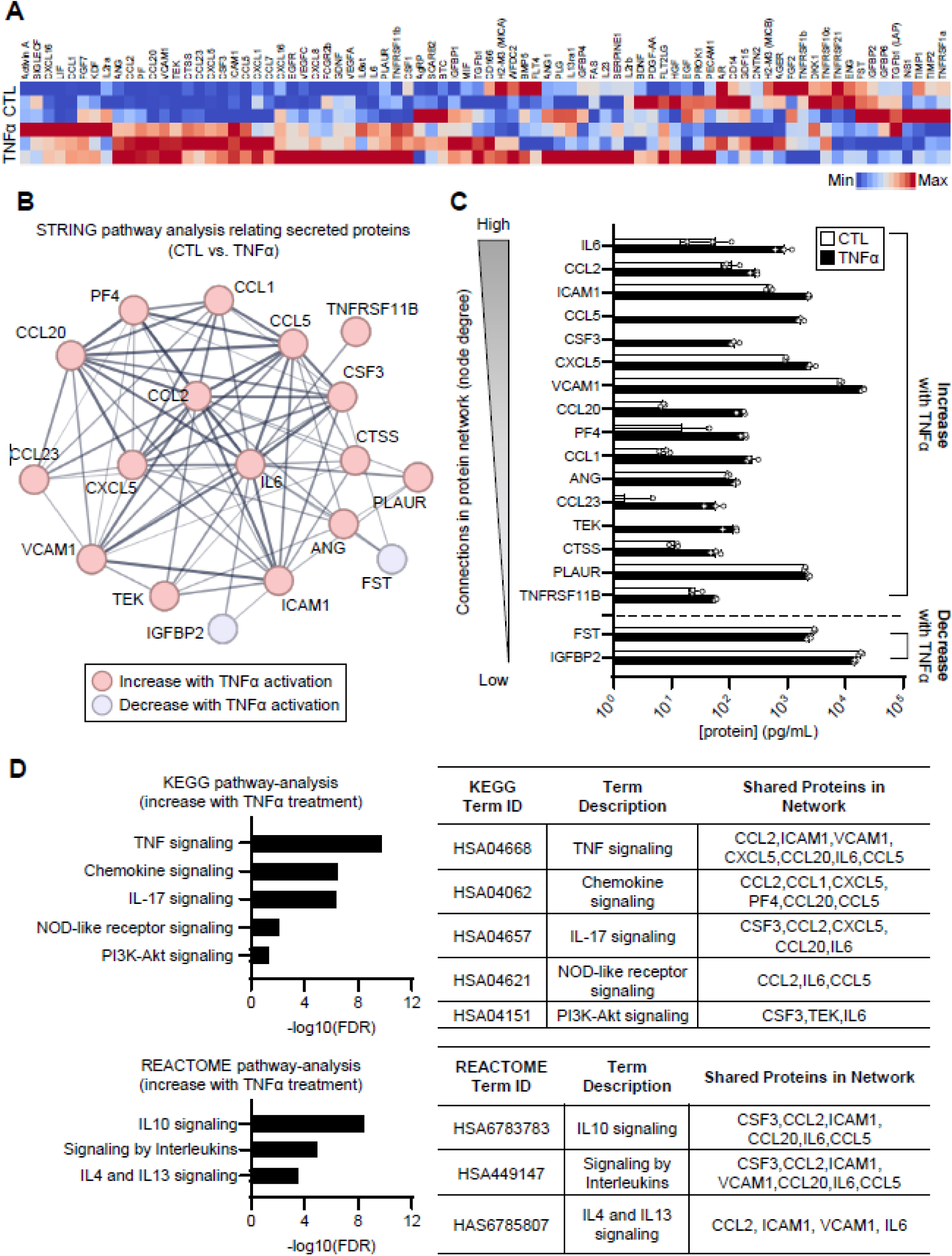
TNF-α stimulation promotes pro-inflammatory secretory phenotype in pericytes. **A.** Heat map representing all the proteins secreted by unstimulated (CTL) and stimulated (TNF-α) pericytes. Two-way clustering performed using Ward method. Red=high secretion, Blue=Low secretion. N=3 replicated conditioned medium samples per condition. **B.** STRING pathway analysis relating secreted proteins. **C.** Quantification of the secreted proteins after stimulation. Proteins are ordered from those with the most connections (node degrees) to the least number of connections within the network, and only the significantly changed proteins (p<0.05) are shown. **D.** Pathway enrichment analysis, KEGG and REACTOME, of the differentially secreted proteins. False discovery rate (log 10) of significant pathways is shown.

To better elucidate the signaling pathways associated with immunological stimulation in pericytes, we performed KEGG and REACTOME pathway analysis on the proteins that significantly increased upon TNF-α treatment (**Figure 2D**). Our analysis identified multiple pathways related to chemokines and interleukins that were upregulated after treatment. IL-17 signaling significantly alters the inflammatory gene expression of pericytes^37^ and has been shown to be a key driver in neuroinflammation^38,39^. NOD-like pattern recognition receptors are expressed in pericytes, and activation of these receptors can lead to the secretion of inflammatory cytokines involved in neuroinflammation^40^. Similarly, PI3K-Akt signaling pathway in pericyte is also associated with extracellular matrix (ECM) degradation and BBB impairment by mediating VEGF expression^41^. These signaling pathways suggested how pericytes could play a mediator role in neuroinflammation. Pathways observed in stimulated pericytes included IL-4/IL-13 signaling, which involved both ICAM1 and VCAM1 in its protein network. Both neuroprotective and neurotoxic effects have been reported regarding IL-4/IL-13 signaling - whether these interleukins reduce inflammation or potentiate oxidative stress during neuroinflammation^42^. Indeed, enhanced expression of IL-4/IL-13 in activated microglia has been shown to play a critical role on neuronal death during Alzheimer’s disease^43^. Additionally, IL-4/IL-13 signaling is characteristic of the T helper type 2 (Th2) immune response, suggesting that an increase in this pathway is linked to eosinophil and other immune cell trafficking^44^. Overall, these data suggest that immunological stimulus of pericytes impacts the expression levels of proteins related to cellular physiology and morphology - such as ICAM1 and VCAM1 - which in turn may initiate downstream immune cascade involved in the regulation of neuroinflammation.

### MSC-EV treatment modulates pericyte morphology

After establishing pericyte morphological and secretory control responses to TNF-α stimulation, we used this morphological assay to assess the effects of MSC-EVs produced under varying conditions. We treated pericytes with 4 different batches of MSC-EVs, generated from two different manufacturing platforms (2D-flasks vs 500 mL vertical-wheel 3D-bioreactors), with or without 50 ng/mL IFN-γ/TNF-α priming (**Figure 3A**). Cytokine priming is a well-established approach to mitigate functional heterogeneity of MSCs and enhance their immunomodulatory function^45^. These manufacturing conditions for MSC-EVs were adopted from a previous study by our group, which demonstrated significant modulation of microglia morphology and their secretome profile^2^.

**Figure 3.**
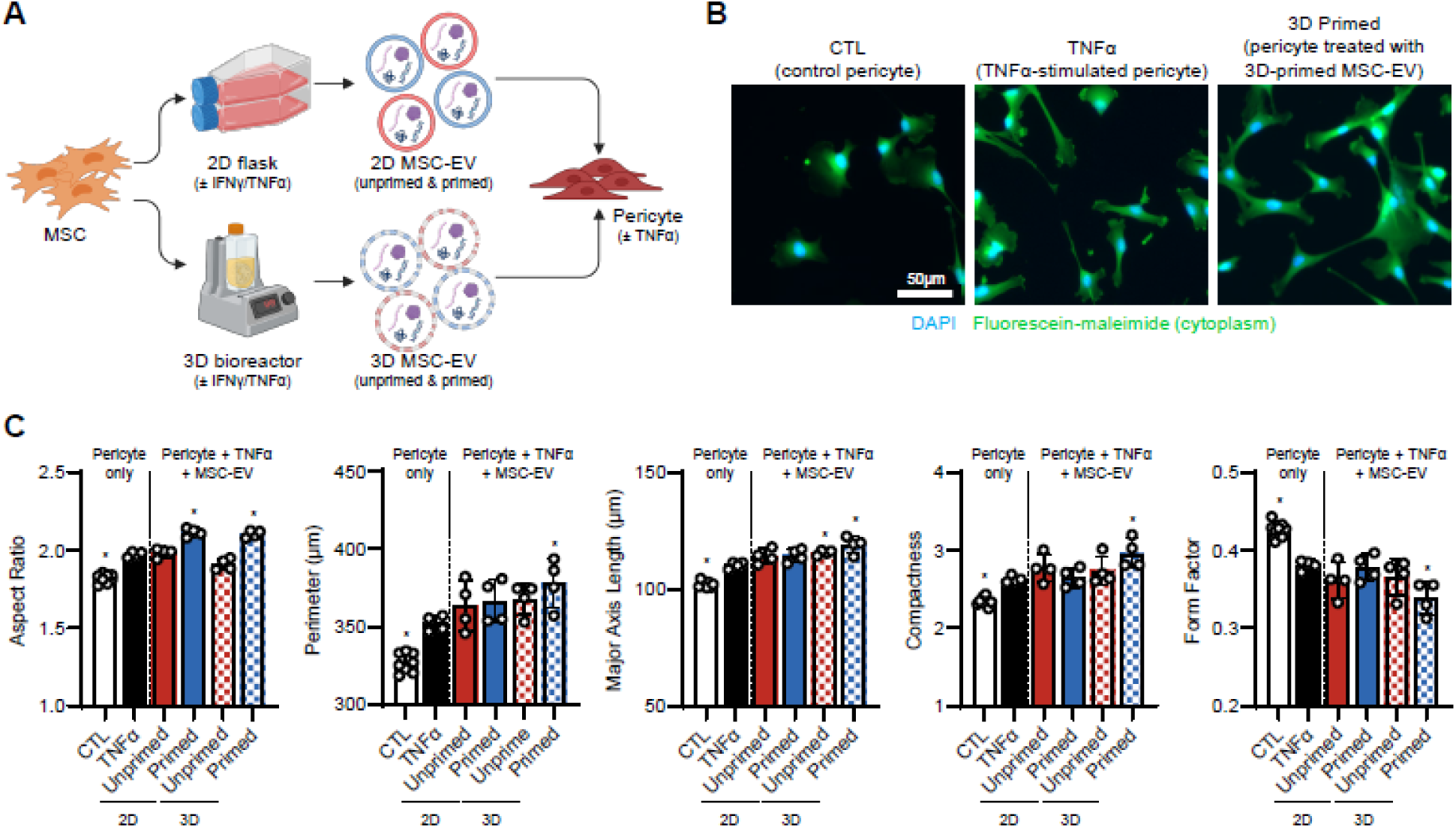
Cytokine priming of MSC-EVs enhances their impact on pericyte morphology. **A**. Schematic illustration of various MSC-EV manufacturing conditions. **B.** Representative images of pericytes treated with different MSC-EVs from various conditions. Pericytes were stained with Hoechst (blue) and fluorescein-maleimide (green) to show nuclei and cytoplasm, respectively. **C.** Quantification of morphological parameters of pericytes in response to various MSC-EV treatment. For each morphological feature, n=6 wells per condition. *p<0.05 relative to the stimulated pericyte group (TNFα). Data plotted as mean ± s.d.

Interestingly, all MSC-EV groups demonstrated a general trend of increased size (perimeter, major axis length) and complexity (compactness, aspect ratio) in the stimulated pericytes—a trend also observed with immunological stimulation alone. Among the MSC-EV groups, those manufactured using IFN-γ/TNF-α priming in a bioreactor exhibited a consistent and significant changes in all morphological parameters, suggesting a robust effect of both the 3D culture platform and priming conditions (**Figure 3B, C**). This highlighted that while TNF-α stimulation alone increased pericyte size, the addition of immunomodulatory MSC-EVs further enhanced this effect, indicating a complex relationship. In fact, this greater morphological change induced by MSC-EVs (i.e., ‘enhancement’ of morphological response to cytokines) in pericytes is contrary to our previous observation in microglia, where MSC-EV treatment resulted in a less pronounced morphological response (i.e. ‘suppression’ of morphological response), suggesting that MSC-EVs promoted an alternate morphological response for pericytes.

### Priming condition and microcarrier concentration impact MSC-EV modulation of pericyte morphology

Changes in pericyte morphology may serve as indicators of underlying functional alterations, and morphology-associated changes can influence critical cell functions, impacting the regulation of inflammation and modulation of blood vessel formation. Therefore, we further explored broader manufacturing and priming conditions for MSC-EVs (**Figure 4**), and profiled secretory phenotypes associated with these morphological changes (**Figure 5**). For future studies, we chose to move forward with the bioreactor manufacturing platform based on several considerations: 1) The bioreactor primed group was the only MSC-EV group that consistently modulated pericyte morphology, and 2) bioreactors are amenable to scaling necessary for clinical production.

**Figure 4.**
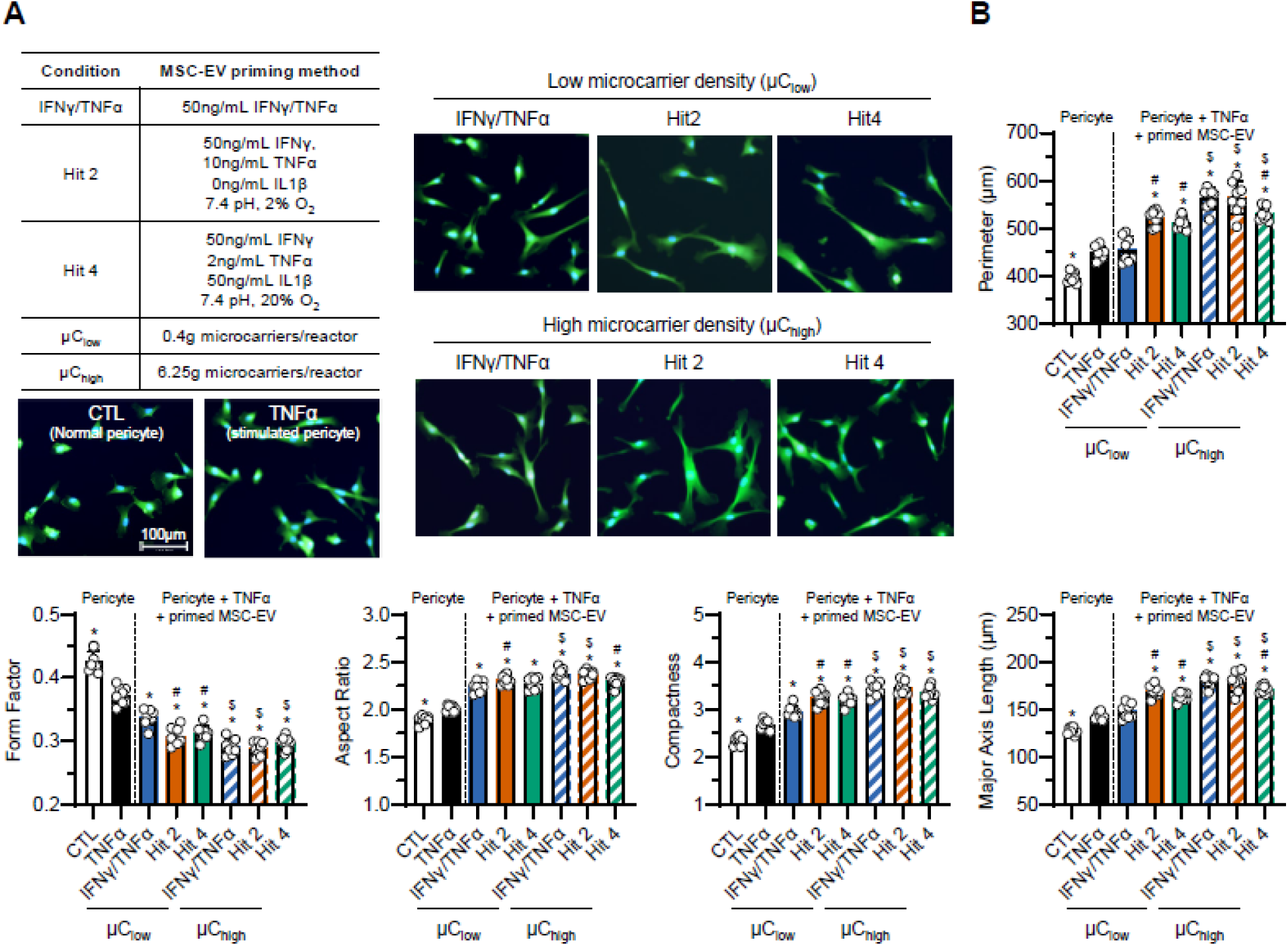
MSC-EV priming conditions and microcarrier density affect pericyte morphology. **A**, Overview of MSC-EV manufacturing conditions (left) and representative images of pericytes cultured under these conditions (right). Pericytes were stained with Hoechst (blue) and fluorescein-maleimide (green) to show nuclei and cytoplasm, respectively. **B,** Quantification of morphological parameters of pericytes treated with MSC-EVs generated from various manufacturing conditions. For each morphological feature, n=6 wells per condition. *p<0.05 relative to the stimulated pericyte group (TNFα). ^#^p<0.05 relative to the IFNγ/TNFα condition within μC density. ^$^p<0.05 relative to μC_low_ within priming conditions.

**Figure 5.**
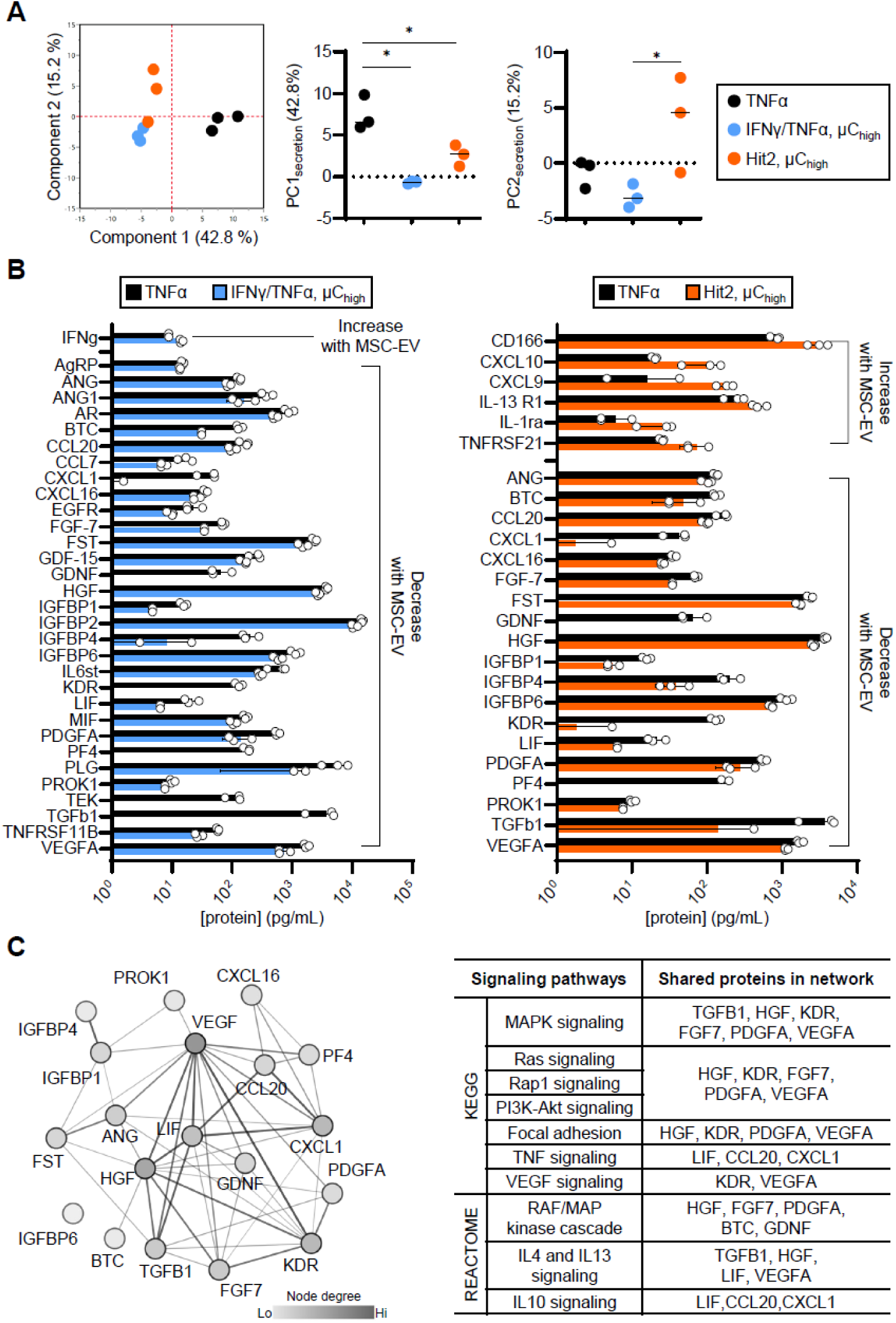
Primed MSC-EVs modulate pericyte secretome. **A,** Individual clusters of TNFα (red), IFNγ/TNFα, μC_high_ (blue), and Hit2, μC_high_ (orange) observed with PCA conducted using all the proteins analyzed. *p<0.05 using one-way ANOVA with Tukey’s multiple comparisons test across all groups. **B,** Quantification of the pericyte proteins affected by MSC-EVs generated from IFNγ/TNFα, μC_high_ priming condition (left) or Hit 2, μC_high_ priming condition (right). **C,** STRING network analysis (left) and KEGG/REACTOME pathway enrichment analysis (right) of downregulated proteins following MSC-EV treatment. The analysis focuses on proteins that were consistently decreased in response to IFNγ/TNFα and μC_high_ MSC-EV, as well as Hit 2, μC_high_ MSC-EV. For all bar graphs, data are mean ± s.d (n=3 replicate conditioned medium sample per group). Only proteins significantly different between EV treatment groups and TNF-α control included as input to PCA in **A, B**.

We explored additional manufacturing conditions that have previously shown more effective cell morphological modulation^2^. MSC-EVs were manufactured in the same 500 mL vertical wheel bioreactor using three different priming conditions and two different microcarrier concentrations with detailed descriptions of each condition tabulated in **Figure 4A**. The IFN-γ/TNF-α priming condition is the same condition shown in **Figure 3A** and the other 2 priming conditions (Hit 2 and Hit 4) were identified as ‘Morphological Hits’ from a high throughput morphological screen from our previous work and shown to have enhanced microglia modulatory activity versus unprimed MSCs^2^.

We assessed the pericyte morphological response to these 6 different MSC-EV preparations using our established analysis pipeline. Similar to our previous observation (**Figure 3**), we observed a notable difference between the TNF-α stimulated pericytes and all the MSC-EV groups, where MSC-EV treatments exhibited much bigger and elongated pericyte morphology (**Figure 4A**). All MSC-EV groups significantly increased complexity (increase in aspect ratio and compactness, decrease in form factor) and size (increase in perimeter and major axis length) compared to the stimulated pericytes except for IFN-γ/TNF-α priming at low microcarrier concentration (µC_low_).

Furthermore, we were able to use pericyte morphology to assess the impact of cytokine priming and microcarrier conditions on MSC-EVs. Analysis of MSC-EV groups manufactured using µC_low_ showed Hit 2 and Hit 4 priming conditions had a significant effect on pericyte morphological response compared to IFN-γ/TNF-α priming (#p<0.05, **Figure 4B**). For MSC-EVs manufactured using high microcarrier (µC_high_) conditions, IFN-γ/TNF-α and Hit 2 priming groups were similar while Hit 4 group had a less significant enhancement effect on pericyte morphology, though still greater than the TNF-α-stimulated control. When comparing the role of microcarrier concentration, we found MSC-EV priming with IFN-γ/TNF-α or Hit 2 condition using µC_high_ enhanced pericyte morphological response more so than those using µC_low_, indicated by all morphological parameters ($p<0.05, **Figure 4B**). For Hit 4 priming conditions, this greater morphological response was only observed in perimeter, major axis length, compactness, and form factor, but not in aspect ratio.

### Primed MSC-EVs modulate secretory phenotype from the stimulated pericytes

We then analyzed how two specific MSC-EV groups, from MSCs primed using IFN-γ/TNF-α or Hit 2 at µC_high_, affected pericyte secretome profile using a panel of 200 chemokines, cytokines and growth factors (**Figure 5**). These 2 groups were chosen as they had the most significant impact on pericyte morphological response to TNF-α (**Figure 4**). All proteins shown in **Figure 5** were detected at levels significantly above pericyte media (and pericyte media + MSC-EV only) controls indicating changes in secretion were from pericytes and not proteins from the MSC-EV prep.

Both IFN-γ/TNF-α and Hit 2 MSC-EV groups largely decreased the secretion of inflammatory cytokines, chemokines and growth factors – such as CXCL1, LIF, TGF β1, and VEGFA. A recent study demonstrated that immunological activation of pericytes significantly upregulated the production of these inflammatory secretome^46^, and MSC-EVs generated from our primed batches were able to significantly regulate their secretion. This is particularly important because many of these secretome can induce pro-inflammatory states not just in pericytes, but also in astrocytes, microglia, and endothelial cells, and further recruit leukocytes to exacerbate neuroinflammation^47,48^. Additionally, the expression of angiogenic factors, such as ANG, BTC, FGF-7, FST, GDNF, HGF, and VEGFA, have all decreased upon MSC-EV treatment in the stimulated pericytes. Neuroinflammation and angiogenesis are closely related process in the progression of CNS disorders, and while angiogenesis may help repair damaged tissue after inflammation, it also is a prominent feature of several CNS diseases^49,50^ as it can actively promote inflammation^51^. Indeed, injection of the well-recognized angiogenesis promoting factor VEGF into rat CNS was shown to induce both angiogenesis and inflammation^52^. Reduction of multiple angiogenic factors by MSC-EVs suggested a potential mechanism of action of our primed MSC-EV therapies in regulating neuroinflammation. Overall, enhanced secretion of immunological secretome after TNF-α stimulation in the pericytes were mitigated by both MSC-EV groups from different conditions.

Furthermore, there were several chemokines (CXCL9, CXCL10) and receptors (IL-1ra, IL-13R1, TNFRSF21, and CD166) that notably increased upon Hit 2 MSC-EV treatment. Both CXCL9 and CXCL10 are chemokines that are known to recruit activated T cells^53^ and regulate vessel stability *via* CXCR3 receptor signaling pathways^54–56^. These chemokines can limit the function of endothelial cells during the resolving phase of wound healing^55,57,58^, highlighting their importance in vascular remodeling during inflammation. Additionally, MSC-EV treatment greatly increased the levels of immunosuppressive membrane molecules IL-1ra^59,60^, IL-13R1^61^, and CD166^62,63^ in pericytes, which all have immunomodulatory roles in T cell suppression, microglia activation, migration of pericytes, and tight junction permeability. Overall, these data underscore the regulatory role of MSC-EVs in modulating immune responses, while highlighting key secretome pathways that may contribute to neuroprotection through their beneficial effects on immune modulation and neural repair.

## DISCUSSION

To fully realize the therapeutic potential of MSC-EVs, and facilitate their translation for neurodegenerative diseases, it is essential to develop robust assays that effectively demonstrate their immunomodulatory activity aligned with their proposed mechanisms of action. Here, we demonstrated how a high-throughput morphological screening bioassay can be applied to assess MSC-EV bioactivity on the target cell phenotype, pericytes. Our findings indicate that pericyte morphological responses are closely linked to the immunomodulatory activity of MSC-EVs across different production batches, particularly in the context of neurodegenerative disease treatment. Using this approach, we screened the effects of different manufacturing conditions on MSC-EV, and demonstrated that MSC-EVs enhanced the morphological response of pericytes, dramatically downregulating their neuroinflammatory secretome (e.g., VEGFA, TGFb1, CXCL1, CXCL16) and upregulating vascular remodeling secretory profile (e.g., CXCL9, CXCL10, IL-1ra, CD166).

Pericyte morphology is closely linked to their physiological roles in the context of neuroinflammation, with different morphologies associated with distinct functionalities^64^. For instance, one study suggested that elongated form of pericytes may increase the gap size between venules to promote neutrophil transmigration^65^. Another study demonstrated that bigger, and more elongated form of pericytes have reduced cellular motility and αSMA expression^66^. In our study, we observed consistent differences in both morphological (**Figure 1**) and secretory phenotype (**Figure 2**) between unstimulated pericytes and those stimulated with TNF-α. The morphological changes following stimulation included increases in cell size and complexity, as reflected by measures in perimeter, major axis length, aspect ratio, compactness, and a reduction in form factor. These results align with previous findings that TNF-α stimulation at 10ng/mL elongated the pericytes phenotype and enhanced their migratory behavior^67^. We believe this morphological change could be reflective of their function *in vivo* where they are known to migrate from sites of injury and hypoxia^68,69^. In addition to morphological changes, TNF-α also influenced pericyte secretion profiles (**Figure 2**). We observed a significant increase in IL-6, ICAM-1 and CCL5 expression in response to TNF-α, mirroring the findings from previous studies^70–72^. IL-6 has been shown to induce BBB dysregulation through the activation of microglia and astrocytes, leading to neuronal damage^70^. ICAM-1 and CCL5 both have roles in recruiting immune cells (i.e., monocytes, macrophages, NK cells, T cells) to the site of inflammation^73,74^. Overall, our findings demonstrate TNF-α, a key mediator of neuroinflammation and several neurodegenerative diseases^32,34^, promotes a pro-inflammatory phenotype in pericytes *in vitro*. This suggests the pericytes may play an active role in recruiting both innate and adaptive immune cells to the site of inflammation *in vivo*, further supporting their involvement in neuroinflammatory processes.

In terms of the effect of EVs on pericyte morphological parameters, we initially hypothesized that EV treatment would suppress the pericyte morphological response to TNF-α (i.e. preventing their increase in size and complexity); however, MSC-EVs rather enhanced the effect in pericyte morphological response to TNF-α, suggesting larger size of pericytes does not necessarily equate to a more inflammatory form. MSC-EVs manufactured using 6 different conditions (**Figure 4A**) increased the pericyte size (increase in perimeter, aspect ratio, and major axis length) and complexity (increase in compactness, and decrease in form factor), while significantly downregulating the secretion of pro-inflammatory secretome from the stimulated pericytes. Interestingly, these changes in cellular morphologies and secretory phenotypes have previously been seen in primed MSCs^27,28^. Cytokine-primed MSCs, treated with either IFN-γ alone or a combination of IFN-γ and TNF-α, exhibited increase in size and complexity, while demonstrating an enhanced immunomodulatory function as seen in their T cell suppression. Although pericytes and MSCs possess different functional properties, they both share similar immunomodulatory activities in *in vitro* models^31,75^. Considering the functional and phenotypic similarities observed between pericytes and MSCs both *in vitro* and *in vivo*, it is notable that they respond similarly to inflammatory signals (e.g. cytokines) in terms of their morphology.

Next, we hypothesized this alternate morphological response is indicative of pericytes having increased immunomodulatory function. We analyzed pericyte secretome in response to MSC-EVs as an indicator of MSC-EV bioactivity and found that MSC-EVs modulated pericyte secretion of proteins related to inflammatory response (**Figure 5**). Among many proteins that decreased with MSC-EV treatment in pericytes, a significant reduction in angiogenic proteins, including ANG, BTC, TGF-β1, and VEGFA, was notable. The role of angiogenesis is context-dependent, and many studies have shown that neoangiogenesis can promote inflammation of the CNS^50,76^. Among these proteins, BTC and VEGFA have shown direct correlation with neuroinflammation. Blocking Epidermal Growth Factor Receptor (EGFR), a receptor for a ligand BTC, can reduce microglial inflammatory response^77^ and improve behavior changes in Alzheimer’s disease^78^. Similarly, a soluble receptor for VEGFA can reduce neuropathic pain development in nerve injury model^79^. Furthermore, our pathway analysis (**Figure 5C**) demonstrated that MSC-EV treatments, regardless of their batch conditions from IFN-γ/TNF-α or Hit 2 priming, downregulated pathways associated with microglial immune response, such as MAPK and PI3K-AKT signaling^80,81^. Collectively, these findings suggest that the primed MSC-EV therapies may indirectly modulate multiple cell-types in the BBB and brain parenchyma with pericytes serving as the key mediator.

MSC-EV manufacturing processes have been shown to change the functional outcomes of the EVs^2,82,83^. In this work, we observed that the magnitude of the enhanced pericyte morphological response was dependent on the manufacturing conditions. Compared to MSC-EVs produced by MSCs cultured in 2D flasks, MSCs cultured in 3D bioreactors produced EVs that consistently enhanced the pericyte morphological response to TNF-α i.e. increased size, elongation, and complexity (**Figure 3**). Bioreactors are better suited for large scale manufacturing of MSC-EVs by enabling production of large, consistent batches of MSC-EVs and potentially mitigating the variability of multiple smaller MSC-EV batches produced using MSCs cultured in 2D flasks (or multilayered cellstacks). In addition, bioreactors allow for a dynamic environment that induces shear stress through stirring of the cell suspension that alters the topographical and surface environment the cells are exposed to, which can influence their behavior such as migration, adhesion, and morphology^84^. In our case, we found that the MSC-EVs produced with higher microcarrier densities had a greater effect on pericyte morphology, with a greater increase in pericyte size and complexity when compared to pericytes treated with MSC-EVs produced by MSCs cultured at lower microcarrier density. While the exact mechanism by which microcarrier density and priming impact MSC-EV bioactivity is unknown, it has been shown that the microcarrier concentration of 1.25g per 90mL medium batch, which is similar to the high microcarrier density we have used in the experiment with 6.25g per 500mL, yielded the highest MSC proliferation than that of the lower (0.75g/90mL) or higher (2.5g/90mL) microcarrier densities^85^. We observed that regardless of priming conditions, MSC-EVs in general exhibited a significant change in pericyte morphology, however, certain conditions (i.e., IFN-γ/TNF-α or Hit 2 priming) induced a greater change than the other (Hit 4 priming). Pericytes treated with IFN-γ/TNF-α MSC-EVs or Hit 2 MSC-EVs significantly secreted proteins that are associated with lipid response, such as CCL2, CXCL6, CXCL8, IL6, TNFRSF1B^2^, and we have previously reported on the differences in the lipidomic profile of these various MSC-EV batches. Hence, further studies looking into batch-specific lipidomic signatures may enlighten the differential mechanisms of batch conditions on the pericyte morphologies.

This is the first study demonstrating MSC-EV bioactivity in the context of modulating pericyte morphology and secretome profile. This novel approach provides a foundation for exploring MSC-EV potential in treating neurodegenerative diseases by using pericytes as an *in vitro* model of MSC-EV bioactivity. In addition, we believe that this assay will aid in standardization efforts that aim to assess MSC-EV bioactivity. Morphology is a rapid, single-cell, high throughout analysis that provides comprehensive information on cellular responses to disease-relevant signals such as inflammatory cytokines. The capabilities afforded by morphological profiling will facilitate standardization of MSC-EV characterization, evaluation of the effects of manufacturing changes on MSC-EV bioactivity, and further contribute to eventual clinical translation. While this study demonstrates the potential of using pericytes as a model target cell for assessing MSC-EV bioactivity relevant to treatment of neuroinflammation, it is important to note that the studies were conducted using pericytes derived from one donor and MSC-EVs produced from a single MSC line. This morphological platform can therefore be applied in the future to evaluate whether pericytes from different donors respond differently to MSC-EVs and whether MSC-EVs from different cell-lines (akin to different manufacturing conditions evaluated in this study) possess different bioactivity on the pericyte morphological response. Additionally, this work provides a foundation for exploring MSC-EV modulation of pericytes in more complex *in vivo* disease models or 3D *in vitro* systems^86,87^ (i.e. microphysiological ‘on-chip’ systems) that could improve our understanding of not only MSC-EV modulation of pericytes, but indirect modulation of other cell-types mediated by pericytes. Finally, this pericyte morphological assay could be used to better understand MSC-EV (or other cell-based therapy) mechanisms of action, the knowledge of which could help further refine MSC-EV manufacturing and facilitate clinical translation.

## Acknowledgements

We thank Drs. Brenton McCright and John Thomas for review of this manuscript.

## Availability of Data and Materials

The data that support these findings are available from the corresponding author upon request.

## Author Contribution Statement

CEC: Methodology, Software, Formal Analysis, Investigation, Writing – Original Draft, AML: Methodology, Formal Analysis, Investigation, KRD: Methodology, Formal Analysis, Investigation, MR: Investigation, SG: Investigation, NG: Conceptualization, Writing – Review and Editing, Supervision, Project Administration, Funding Acquisition JH: Conceptualization, Formal Analysis, Writing – Original Draft, Writing – Review and Editing, Project Administration, RAM: Conceptualization, Methodology, Formal Analysis, Writing – Original Draft, Writing – Review and Editing, Supervision, Project Administration, Funding Acquisition.

