## supplemental figures and tables for "Development of a High-throughput Morphological Assay for Evaluating Mesenchymal Stromal Cell-derived Extracellular Vesicle Modulation of Brain Pericyte Secretory Phenotype"

| **Measurements made by CellProfiler** | **Description** |
|---|---|
| Perimeter | The total number of pixels around the boundary of each region in the image. |
| Major Axis Length | The length of the major axis of the ellipse that has the same normalized second central moments as the region. |
| Compactness | Calculated as $Perimeter^2/4*\pi*Area$, related to Form Factor. A filled circle will have a compactness of 1, with irregular objects or objects with holes having a value greater than 1. |
| Form Factor | Calculated as $4*\pi*Area/Perimeter^2$. Equals 1 for a perfectly circular object. |
| Aspect Ratio | Major axis length divided by minor axis length. |

**Supplementary Table 1.** Table defining 5 morphological features quantified in the study using CellProfiler.

| Proteins detected as secreted by pericytes at levels above the pericyte medium-only control | | | | | | | | |
|---|---|---|---|---|---|---|---|---|
| Activin A | BMP-5 | EGF | CSF3 | IGFBP1 | CXCL8 | MICB | PF4 | TNF RII |
| AgRP | BTC | EGF R | CXCL6 | IGFBP2 | INS | MIF | MOK | TRAIL R3 |
| ALCAM | CTSS | CXCL5 | GDF15 | IGFBP4 | LAP(TGFb1) | CCL20 | CCL5 | PI3 |
| ANG1 | CD14 | ENG | GDNF | IGFBP6 | LIF | CCL23 | Siglec-5 | PLAUR |
| ANG | CNTN2 | FAS | LRPPRC | IL-13 R1 | SCARB2 | NTF4 | TGFb1 | VCAM-1 |
| PLG | CXCL16 | Fcg RIIBC | CXCL1 | IL-2 Ra | CCL2 | TNFRSF11B | TEK | VEGF |
| AR | DKK-1 | FGF-7 | HGF | IL-2 Rb | CCL7 | SERPINE1 | TIMP-1 | VEGF R2 |
| BDNF | TNFRSF21 | Flt-3L | CCL1 | IL-23 | CSF1 | PDGF-AA | TIMP-2 | VEGF R3 |
| bFGF | PROK1 | FST | ICAM-1 | IL-6 | MICA | PECAM-1 | TNF RI | VEGF-C |

**Supplementary Table 2.** Comprehensive list of proteins secreted by pericytes detected above levels in pericyte medium-only control.

| Protein | Identifier | Node degree | CTL_1 | CTL_2 | CTL_3 | TNF-α_1 | TNF-α_2 | TNF-α_3 |
|---|---|---|---|---|---|---|---|---|
| IL6 | 9606.ENSP00000385675 | 17 | 51.3 | 108.8 | 18.5 | 869.0 | 617.7 | 1,205.6 |
| CCL2 | 9606.ENSP00000225831 | 14 | 83.5 | 151.8 | 88.9 | 227.4 | 300.5 | 304.7 |
| ICAM1 | 9606.ENSP00000264832 | 12 | 501.8 | 427.8 | 553.1 | 2,438.2 | 2,303.9 | 2,340.6 |
| CCL5 | 9606.ENSP00000474412 | 11 | 0.0 | 0.0 | 0.0 | 1,694.8 | 1,946.0 | 1,545.5 |
| CSF3 | 9606.ENSP00000225474 | 11 | 0.0 | 0.0 | 0.0 | 107.3 | 148.6 | 118.5 |
| CXCL5 | 9606.ENSP00000296027 | 11 | 983.0 | 917.5 | 934.5 | 2,323.7 | 3,099.7 | 2,419.8 |
| VCAM1 | 9606.ENSP00000294728 | 11 | 7,895.7 | 8,900.0 | 8,055.6 | 18,642.6 | 20,989.2 | 21,301.6 |
| CCL20 | 9606.ENSP00000351671 | 10 | 7.8 | 6.5 | 7.3 | 131.4 | 185.0 | 176.2 |
| PF4 | 9606.ENSP00000296029 | 10 | 0.0 | 45.0 | 0.0 | 154.8 | 195.2 | 192.8 |
| CCL1 | 9606.ENSP00000225842 | 8 | 8.2 | 9.5 | 6.1 | 314.0 | 203.2 | 218.2 |
| ANG | 9606.ENSP00000336762 | 7 | 93.3 | 100.8 | 90.5 | 121.1 | 135.2 | 139.6 |
| CCL23 | 9606.ENSP00000481357 | 7 | 0.0 | 4.8 | 0.0 | 57.5 | 79.2 | 36.3 |
| TEK | 9606.ENSP00000369375 | 6 | 0.0 | 0.0 | 0.0 | 76.5 | 130.5 | 134.6 |
| CTSS | 9606.ENSP00000357981 | 5 | 12.5 | 12.9 | 9.1 | 58.9 | 70.7 | 47.4 |
| PLAUR | 9606.ENSP00000339328 | 4 | 2,066.5 | 1,924.2 | 2,043.6 | 2,276.6 | 2,286.7 | 2,457.2 |
| TNFRSF11B | 9606.ENSP00000297350 | 1 | 22.5 | 23.1 | 34.1 | 58.2 | 55.3 | 60.2 |
| FST | 9606.ENSP00000256759 | 3 | 2,840.2 | 3,045.0 | 2,925.6 | 2,426.1 | 2,141.8 | 2,542.8 |
| IGFBP2 | 9606.ENSP00000233809 | 2 | 17,001.1 | 20,017.9 | 18,614.4 | 15,231.4 | 14,005.5 | 14,477.5 |

**Supplementary Table 3.** Comprehensive list of upregulated (orange) and downregulated (grey) proteins after TNF-α stimulation in pericytes. Proteins are ordered by the node degree.
